# *Listeria monocytogenes* personalized cancer vaccines drive therapeutic immune responses to cancer derived neoantigens

**DOI:** 10.1101/2020.05.11.088930

**Authors:** Brandon Coder, Olga Pryshchep, Dipti Kelkar, Elena Filippova, Xiaoming Ju, David Balli, Cristina Mottershead, Kim Ramos, Nithya Thambi, Zhiyong Cheng, Bryan Vander Lugt, Justin Lesch, Xian Liu, Jason DeVoss, Keegan Cooke, Siyuan Liu, Jinghui Zhan, Petia Mitchell, Daniel O Villarreal, Sandra M. Hayes, James A Johnston, Robert Petit, Hyewon Phee, Michael F. Princiotta

## Abstract

**Background:** Recent advances in the field of cancer immunotherapy have identified CD8^+^ T cell responses against tumor-specific mutations as a key driver of tumor regression and overall survival. ADXS-NEO is a personalized *Listeria monocytogenes* (*Lm*)-based immunotherapy designed to target a patient’s mutation-derived tumor-specific neoantigens. The objective of this study is to demonstrate the feasibility of using the ADXS-NEO platform to target tumor-specific point mutations and control tumor growth by generating neoantigen-specific T cell responses using a pre-clinical mouse tumor model.

**Methods:** Whole-exome sequencing of the MC38 mouse tumor cell line identified 2870 unique non-synonymous mutations. The netMHCcons algorithm was used to predict 137 potential neoantigens. We validated 20 immunogenic neoantigens either by peptide immunization followed by ELISPOT or by the presence of CD8^+^ T cells recognizing the neoantigen peptide following checkpoint inhibitor treatment. Two ADXS-NEO vectors were constructed; Lm20, targeting 20 validated immunogenic neoantigens, and Lm19, targeting most of the non-validated NSMs.

**Results:** Both Lm19 & Lm20 significantly slowed tumor growth in C57BL/6 mice compared to control. An accumulation of ADXS-NEO-specific TILs was observed in tumor bearing mice treated with either Lm19 or Lm20. Examination of the tumor microenvironment in Lm19 or Lm20 treated mice revealed a decrease in the frequency and absolute number of Tregs, TAMs, MDSCs, and PD1^high^ exhausted CD8^+^ T cells as well as an increase in the frequency and absolute number of effector CD8^+^ T cells, relative to control.

**Conclusion:** ADXS-NEO is a potent immunotherapy capable of driving immune responses against tumor-specific mutations and leading to tumor control in mice.

## Background

Genetic alterations found in cancer cells, but not in non-malignant cells, that result in non-synonymous somatic mutations (NSM) can drive anti-tumor T cell responses^1–5^. Cancer-reactive CD8^+^ or CD4^+^ T cells can recognize mutated (MT) epitopes (neoepitopes) from neoantigens presented in the context of major histocompatibility complex (MHC) class I or class II molecules, respectively^6^. CD8^+^ T cells recognizing immunogenic neoepitopes are activated and proliferate, but ultimately become exhausted and express high levels of PD-1 as a result of prolonged exposure to antigen ^7,8^. Anti-PD-1 or anti-PD-L1 checkpoint inhibitor cancer immunotherapy aims to reverse this exhaustion by blocking the activity of checkpoint molecules such as PD-1^9^. Indeed, recent clinical studies using checkpoint inhibitor therapy demonstrated that neoantigen burden is a critical and predictive biomarker for responses to checkpoint therapy^1,10-12^, suggesting T cell responses to neoantigens may control cancer growth if checkpoint inhibitors are provided. However, checkpoint inhibitor trials have demonstrated limited clinical efficacy as a monotherapy^14^ suggesting combination with other immunotherapies will be necessary for optimal benefit.

While checkpoint inhibition may help expand pre-existing populations of T cells, responses mediated by newly generated T cells require priming to the appropriate tumor-specific target antigens in the correct immunologic context^6,14,15^. One method of eliciting *de novo* T cell responses is through immunization using therapeutic cancer vaccines, which have been in development over the past two decades but have had limited success^17^. However, recent Phase 1 clinical studies using neoantigen-targeting cancer vaccines to treat advanced stage melanoma have re-established the potential of therapeutic vaccines for use in immunotherapy^17–21^. The recent success of adoptive T cell therapies targeting patient-specific neoantigens supports this approach^23^.

There are many outstanding questions and challenges to overcome before cancer vaccines become a viable therapy. First, the anti-tumor activity of neoantigen-specific CD8^+^ vs CD4^+^ T cells needs to be clarified. Interestingly, two previous cancer vaccine studies reported a strong CD4^+^ T cell bias when either peptide or mRNA was used for antigen delivery^19,20,23^. While the roles of CD4^+^ T cells as providing help to CD8^+^ T cells for differentiation into memory cells by providing IL-2, and, also as contributing directly as cytotoxic effector CD4^+^ T cells^24^, designing cancer vaccines to generate CD4^+^ T cells targeting MHC class II epitopes would not seem to be the most efficient approach because with the inherent cytotoxic ability of CD8^+^ T cells to kill tumor cells. In addition, there is currently no algorithm that can reliably predict the immunogenicity of a neoepitope ^25,26^. As such, the capacity of a cancer vaccine to deliver multiple antigenic targets becomes far more critical, as the more neoantigens the vaccine targets, the greater the probability of eliciting a neoantigen-specific T cell response.

We initiated a study to test the feasibility of using a bacterial vector-based cancer vaccine as an immunotherapy targeting tumor derived neoantigens. We evaluated the potential of using *Listeria monocytogenes* (*Lm*) as a personalized cancer vaccine (PCV), known as ADXS-NEO, utilizing the highly attenuated LmddA strain^27^. LmddA-based vectors are bioengineered to secrete one or more tumor-associated antigens fused to a tLLO peptide, including the tLLO secretion signal, transcribed from an extrachromosomal bacterial plasmid ^28^. The LmddA strain is attenuated by the irreversible deletion of the *actA* virulence factor, which inhibits bacterial spreading^29^. *Lm*-based vectors are rapidly phagocytosed by host antigen presenting cells (APCs)^31^, which results in the activation and maturation of not only the APC but also other innate immune cells, including neutrophils, macrophages, natural killer (NK) cells, and γδ T cells^30^. Activation of innate immune cells is facilitated by the recognition of *Lm*-specific PAMPs (pathogen-associated molecular patterns) and DAMPs (damage-associated molecular patterns), including TLR2, TLR9, TLR5, and activation of the STING pathway^30,31^. Additionally, *Lm* infection drives a robust CD8^+^ T cell response and establishment of CD4^+^ T cell dependent T cell memory^31^. Once inside the APC, the *Lm*-based vector escapes the phagosome and enters the host cell cytosol, where it secretes the tLLO fusion proteins containing the targeted TAAs^30^, which are then available to the MHC class I antigen processing and presentation pathway. Bacteria that do not escape the phagosome will ultimately be degraded in the phagolysosome, where antigenic proteins enter the MHC class II antigen presentation pathway^32^. Simply stated, the live attenuated bacterial vector acts as a unique self-enclosed antigen delivery system, expressing tumor antigens in the context of the initial stages of a bacterial infection with multiple strong adjuvant properties that allows for the simultaneous activation of innate and adaptive immunity.

Here, we report the use of *L. monocytogenes* to target tumor-derived neoantigens. Using a pre-clinical cancer model, we demonstrate a potential PCV pipeline starting with patient tumor biopsy, whole-exome sequencing, neoantigen prediction and engineering and production of a baceterial vector (ADXS-NEO) with the capacity to target upwards of 40 putative neoantigens per individual construct. We found that ADXS-NEO promotes a pro-inflammatory tumor microenvironment (TME), attenuates intratumoral immune suppression, and drives a robust neoantigen-specific cytotoxic T lymphocytes (CTL) response. This targeted inflammation inhibits tumor progression, extends overall survival, and confers long-lasting protective immunity.

## Methods

### Comparative Whole Exome Sequencing (WES), RNA-seq, and data analysis

#### Next-generation sequencing

Whole-exome sequencing libraries were constructed using the SureSelect Mouse all Exon Kit (Agilent). RNA sequencing libraries were constructed using the TrueSeq Stranded mRNA Sample Prep kit (Illumina). All sequencing was done on the Illumina HiSeqX platform.

#### Whole-exome and RNA sequencing, analysis and neoantigen identification

Paired-end reads were aligned to the mouse reference genome (GRCm38) using BWA-MEM aligner (version 0.7.15-r1140). Duplicate reads were removed from sorted alignment maps using Sentieon Dedup followed by indel realignment around known indels using Sentieon Realigner and base recalibration using Sentieon QualCal. Somatic variant analysis was performed using Sentieon TNseq Haplotyper algorithm (version 201711.02). Identified variants were annotated using Snpeff (version 4.3p). Alignment to the mouse reference genome (GRCm38) was performed using STAR aligner (version 2.4.1). Expression was determined using HTSeq (version 0.6.1). Read count data were normalized using DESeq2 (version 1.22.1). Somatic missense mutations were filtered for RNA expression (TPM ≥ 0.5) and potential MHC class I peptides with predicted binding affinities for mouse H-2 K^b^ and H-2 D^b^ alleles ≤ 500 nM were considered potential neoantigens. All sequencing and analysis performed at MedGenome.

### Mice and animal care

#### Animal studies were performed at Amgen and Advaxis

##### Amgen

All experimental studies performed at Amgen were conducted under protocols approved by the Institutional Animal Care and Use Committee of Amgen. Animals were housed at Association for Assessment and Accreditation of Laboratory Animal Care International-accredited facilities (at Amgen) in ventilated micro-isolator housing on corncob bedding. Animals had access *ad libitum* to sterile pelleted food and reverse osmosis-purified water and were maintained on a 12:12 hour light: dark cycle with access to environmental enrichment opportunities. Female C57BL/6 mice (Charles River Laboratories), 6 to 8 weeks of age were cared for in accordance with the “*Guide for the Care and Use of Laboratory Animals*, 8th Edition”.

##### Advaxis

Female 8-12-week-old C57BL/6 (Jackson Labs & Charles River Labs) were used in compliance with protocols approved by the Institutional Animal Care and Use committee of Advaxis, Inc. and in accordance with guidelines from the National Institutes of Health.

### Cell line

MC38 cells were maintained in accordance with the Kerafast MC38 cell maintenance protocol. Cells were tested for mycoplasma every 12 months.

### Reagents

SIINFEKL peptide (VACSIN) and CpG(tlr-1826-1) were from InvivoGen (San Diego, CA). Neoantigen peptides were purchased from LifeTein (Somerset, NJ) and BioSynthesis (Lewisville, TX). Dextramers were purchased from Immudex (Copenhagen, Denmark).

### Peptide immunization in naive mice

Mice were immunized twice at a 7 day interval with either 9-mer or synthetic long peptides mixed with CpG intravenously (100 µg peptide with 50 µg CpG in 200 µl PBS per mouse). Splenocytes were harvested 7 days following the second immunization for immunological analysis. Amino acid sequences of the peptides used for immunization are shown in online supplementary table 1. The sequences of three NeoAg synthetic long peptides used for the efficacy study (Fig.1e) are; Adpgk (GIPVHLELASMTNMELMSSIVHQQVFPT), Dpagt1 (EAGQSLVISASIIVFNLLELEGDYR), and Reps1 (GRVLELFRAAQLANDVVLQIMELCGATR). The sequence of the 27-mer MT SLP of Adpgk is TGIPVHLELASMTNMELMSSIVHQQVF.

**Figure 1.**
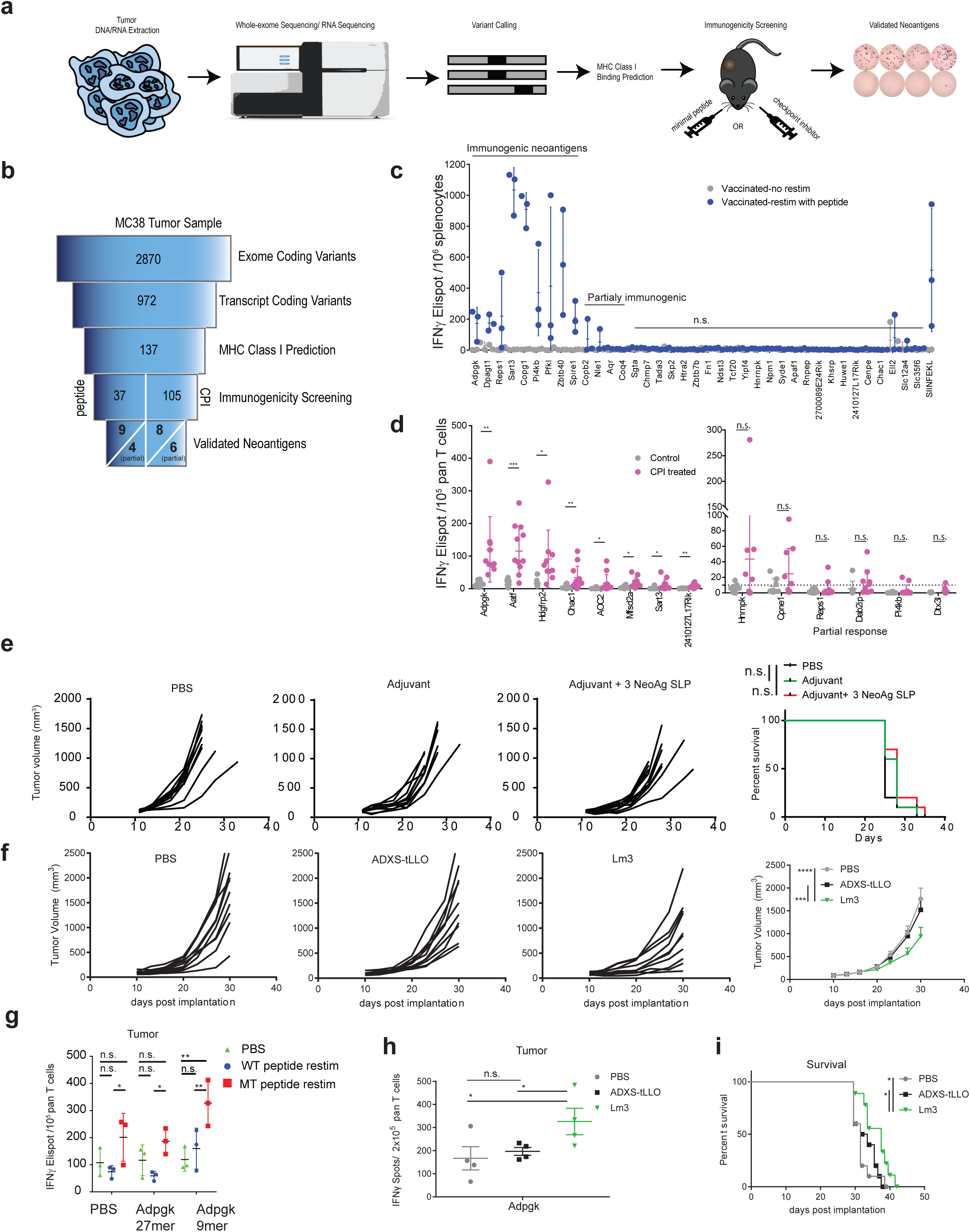
Immunogenicity screening of neoantigens from MC38 cancer cell line. **a**, Illustration demonstrating the process of the neoantigen calling and validation. **b**, Flowcharts of neoantigens identified at each stage of analysis and the number of neoantigens tested in immunogenicity screening and validated neoantigens. **c**, IFNγ Elispot analysis using cells from spleens of mice following immunization of minimal mutated (MT) peptide (9mer peptide harboring mutated amino acid) (n=3 per peptide), re-stimulated in the presence or absence of the minimal MT peptide. **d**, IFNγ Elispot analysis re-stimulated with minimal MT peptides using splenic cells from MC38-implanted mice treated with Control or anti-CTLA4 and anti-PD-L1 (CPI treated). Shown are IFNγ Elispot counts from re-stimulated conditions with minimal MT peptides after substracting the counts from the no peptide control. *Left*; Neoantigens showing significant differences in IFNγ Elispot count between control and CPI treated; *Right*: Neoantigens show in some mice but not statistically significant due to variability of the IFNγ Elispot count. **e**, Tumor volume measurement and survival study of the MC38-transplanted mice treated with either PBS, adjuvant (CpG plus anti-CD40) alone, and adjuvant (CpG plus anti-CD40) plus three synthetic long peptides encoding Adpgk, Dpagt1 and Reps1. **f**,**i**, Tumor volume measurement (**f**) and survival study (**i**) of the MC38-transplanted mice treated with ADXS-tLLO (ADXS-NEO expressing tLLO of listeriolysin, empty vector control) or Lm3 (ADXS-NEO expressing SLP sequences of Adpgk, Dpagt1 and Reps1 as a fusion protein of tLLO). **g**, IFNγ Elispot analysis of pan T cells from the MC38 tumor immunized with synthetic long MT peptide (27mer) or minimal peptides (9mer) encoding Adpgk MT sequences. Re-stimulation was performed using Adpgk 9mer MT peptide. **h**, IFNγ Elispot analysis of pan-T cells from the MC38 tumor vaccinated with ADXS-tLLO or Lm3. *Statistics*, **c-d**, Unpaired t-test, *, 0.01< P <0.05; **, 0.001< P < 0.01; ***, 0.0001< P <0.001; ****, P < 0.0001., n.s., not significant, **f**, Two way ANOVA and Tukey multiple comparison, ***, 0.0001< P <0.001; ****, P < 0.0001., **g**, Two way ANOVA for multiple comparison, *, 0.01< Adjusted P <0.05; **, 0.001< Adjusted P < 0.01. **h**, Unpaired t-test, *, 0.01< P <0.05, **i**, Survival analysis Log-rank test *, 0.01< P <0.05.

### Screening of T cell responses recognizing minimal MHC class I binding neoantigen peptides derived from neoantigens in checkpoint inhibitor-treated mice

MC38 cells were injected subcutaneously on the right flank of mice (3×10^5^ cells) on study day 0. Tumor volume (mm^3^) was measured twice per week using electronic calipers. Once the tumors reached an average of approximately 50 mm^3^ (study day 7), animals were randomized into treatment groups (25 mice per group). Animals were then administered three intraperitoneal injections of anti-CTLA-4 (30 μg/mouse) and anti-PD-L1 (200 μg/mouse) or vehicle every third day. On day 15 spleens were harvested and pan-T cells were isolated as described and stored in liquid nitrogen. Screening to detect T cells recognizing minimal peptides from MC38 neoantigens was performed using a peptide restimulation mouse IFNγ ELISpot assay (Cellular Technology Limited). Briefly, 1×10^5^ pan-T cells were mixed with 3×10^5^ antigen presenting cells (CD3-depleted splenocytes from naïve C57BL/6 mice) along with the minimal neoantigen peptides at a final concentration of 1 µmol/L. The assay was incubated at 37 ° C for 18 hours and IFNγ+ spots were enumerated using a Fluorospot analyzer (Cellular Technology Limited). Anti-CTLA-4 (clone 9D9, mIgG2a) and anti-PD-L1 (clone MIH5, mIgG1) were produced internally at Amgen, Inc.

### Peptide immunization of MC38-implanted mice

Six to eight-week-old C57BL/6 mice were implanted subcutaneously with MC38 tumor cells (3 x 10^5^ cells per mouse). When tumors were palpable (50-150 mm^3^), mice were randomized and peptides with CpG (or PBS control) were administered intravenously, followed by booster immunization seven days later and tumor size was measured and spleens and tumors were harvested seven days following final immunization.

### IFNγ ELISpot assays of splenocytes from immunized mice

IFNγ ELISpot assays were performed by coating plates overnight with antibody, washing with cRPMI once and blocking with 200 µl of cRPMI for 2 hours at room temperature. Half a million to one million splenocytes, or 1 x 10^5^ pan-T cells from tumors, were plated per well in 100 µl of cRPMI. For splenocyte stimulation, 100 µl of peptide was added at a final concentration of 10 µg/ml. For pan-T stimulation, 3 x 10^5^ antigen presenting cells (CD3-depleted splenocytes from naïve C57BL/6 mice) were added per well. Cells were incubated at 37 °C, and IFNγ production was determined after 18 hrs following manufacturer’s instructions (551083, BD Biosciences).

### Dextramer staining

For dextramer staining, a total of 1 x 10^6^ splenocytes or 2 x 10^6^ tumor cells were washed twice with FACS buffer (PBS, 2% FBS, 2mM EDTA), stained with dextramers (Immudex) for 10 minutes at room temperature. Without additional washing, an antibody cocktail containing Fc block (14-0161-86, eBioscience) BV buffer (566349, BD Bioscience) were added. Cells were incubated at 4°C for 20 minutes, washed in FACS buffer, stained with fixable viability dye (65-0866-14, eBioscience) for 10 minutes at room temperature, washed with FACS buffer and flow cytometry was performed within 2 hours of final wash.

### Tumor re-challenge experiment

For tumor re-challenge experiment, mice that had cleared tumor following vaccination and remained tumor-free ≥100 days after tumor implantation and naïve age-matched control mice were implanted subcutaneously with 3 x 10^5^ MC38 tumor cells. Tumors grew to palpable size in all naïve mice after 10 days, while mice that had cleared tumor did not show any sign of tumor growth. Spleens were harvested from all mice and immunophenotype analysis and ELISpot assay were performed.

### Flow cytometry and antibodies

Flow cytometric analyses were done at either the Amgen Flow Cytometry Core Facility (South San Francisco) using an LSRII (BD) flow cytometer, or at Advaxis Inc. using an Attune Nxt (Thermo Fischer) flow cytometer. For surface staining, cells were incubated with fluorochrome-conjugated antibodies for 30 minutes at 4°C using 1:200 dilutions of each antibody (unless otherwise specified). Cells were washed twice in FACS buffer prior to analysis or intracellular staining using the Foxp3/Transcription Factor Staining Buffer Set (eBioscience). Dead cells were excluded either using DAPI (Life technologies) or LIVE/DEAD® Fixable Dead Cell Stain Kits (Life technologies) or fixable viability dye (65-0866-14, eBioscience). Forward and side scatter was used to identify live lymphocytes. Data analysis was performed using FlowJo (version 9.6.2) software (Tree Star).

### ADXS-NEO vector design

Target sequences utilized in vector design are listed in online supplementary table 1. Briefly, the point mutation amino acid was flanked by 7-13 wildtype amino acids, e.g. XXXXXXXXXXMXXXXXXXXXX, where X is the wildtype sequence and M is the mutant amino acid. Total length of the target sequence per NSM was 21 amino acids, except that the NSM was localized towards to the C-terminal end of the protein.

### Plasmid construction, DNA synthesis, ligation, subcloning, transformation, and vaccine production

Target amino acid sequences were reverse-translated, and codon optimized for *Listeria monocytogenes* using OPTMIZER (genomes.urv.es/OPTIMIZER/), and DNA was synthesized. Insert DNA was ligated into the pUC57 shuttle vector and subsequently digested and ligated into the pAdv134 plasmid. ADXS-NEO plasmids were then separately transformed into electrocompetent *L. monocytogenes* strain LmddA to generate strains Lm3, Lm19, Lm20, and Lm19+20. The presence of the correct plasmid within each strain was verified by colony PCR specific for each NEO plasmid (data not shown).

### Immunization of MC38-implnated mice with Lm19 and Lm20

ADXS-NEO vaccine was thawed from -80°C in a 37°C heat block, pelleted at 14,000 rpm for 2 minutes and supernatant was discarded. The pellet was then washed 2X with PBS and re-suspended in PBS to a final concentration of 1.5×10^8^ CFU/mL. MC38 tumor bearing mice were dosed intravenously with 200 μL per mouse using a 29G syringe (BD #324704) for a total of 3 x 10^7^ CFU per mouse, once per week starting at day 10 post tumor implantation for a total of 3 doses, unless otherwise specified.

## Results

### Neoantigen identification in the MC38 tumor model

The aim of this study was to test the feasibility of a comprehensive personalized cancer vaccine targeting NSMs identified using next-generation sequencing. We performed comparative whole-exome sequencing of the murine MC38 tumor cell line and C57BL/6 normal tissue. Out of 2870 NSMs identified, NSMs were prioritized based on RNA expression by selecting mutations with transcripts per million (TPM) ≥ 0.5 based on RNA sequencing, which yielded a total of 972 NSMs (Fig. 1a-b). The NSMs were then filtered by predicted MHC class I binding affinity using the NetMHCcons algorithm. A total of 137 NSMs were predicted to bind H-2 K^b^ or H-2 D^b^ with an IC50 ≤ 500 nM and were considered potential neoantigens. Next, we screened the potential neoantigens predicted to bind MHC class I for their immunogenicity using two methods (Fig. 1a-d, Fig. 2a). As a first strategy, C57BL/6 mice were immunized with minimal peptides (9mer MT peptide) containing the mutated NSM and CpG as adjuvant. Seven days later, spleens were harvested from immunized mice and the presence of neoantigen-specific CD8^+^ T cells were evaluated by IFNγ ELISpot. We screened 37 NSMs and detected neoantigen specific CD8^+^ T cell responses against 13 of these NSMs (Fig. 1c, online supplementary figure 1a-b, and online supplementary table 1).

**Figure 2.**
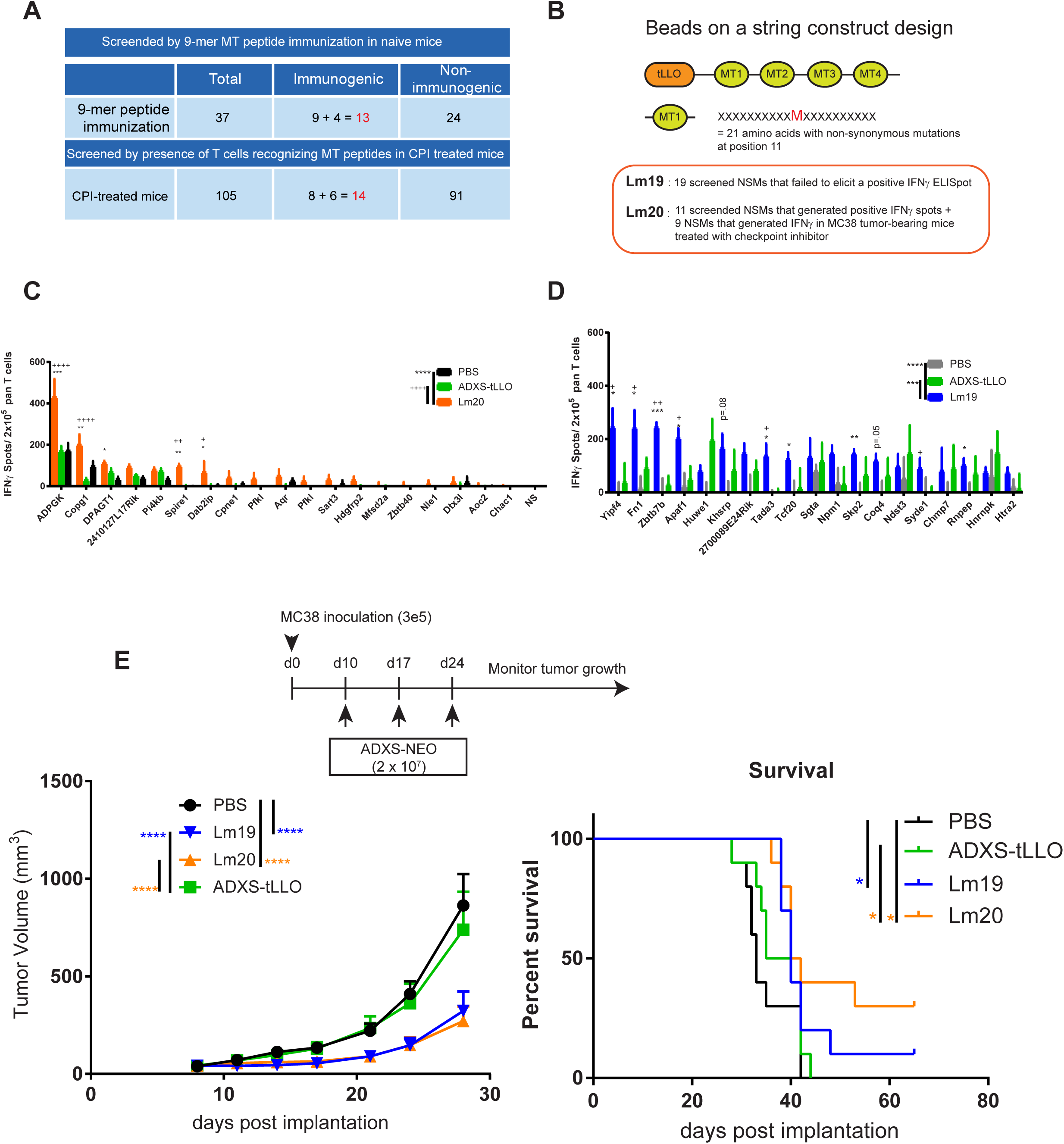
Immunogenicity and efficacy of Lm19 and Lm20 in MC38 tumor model. **a**, Numbers of neoantigens tested in immunogenicity screening *in vivo* and validated neoantigens. **b**, Illustration demonstrating the construct design of Lm19 and Lm20. **c**, IFNγ Elispot analysis from tumors of the mice immunized with Lm20. **d**, IFNγ Elispot analysis from tumors of the mice immunized with Lm19. **e**, Efficacy of Lm19 and Lm20 vaccination in MC38 model. *Statistics*, **c-d**, Unpaired t-test, *, 0.01< P <0.05; **, 0.001< P < 0.01; ***, 0.0001< P <0.001; ****, P < 0.0001., * compares to PBS, + compares to ADXS-tLLO, Two way ANOVA for treatment comparison*, 0.01< P <0.05; **, 0.001< P < 0.01; ***, 0.0001< P <0.001; ****, P < 0.0001. **e**, Two way ANOVA and Tukey multiple comparison *, 0.01< P <0.05; **, 0.001< P < 0.01; ***, 0.0001< P <0.001; ****, P < 0.0001.; Survival analysis Log-rank test *, 0.01< P <0.05; **, 0.001< P < 0.01; ***, 0.0001< P <0.001; ****, P < 0.0001.

As a second strategy to identify immunogenic NSMs, we utilized the fact that checkpoint inhibitor treatment can lead to the activation and proliferation of neoantigen-specific T cells. Therefore, we treated MC38 tumor bearing mice with anti-CTLA4 and anti-PD-L1 antibodies (CPI) and screened minimal peptides derived from 105 predicted NSMs. We identified 8 NSMs capable of elevating CD8^+^ T cell responses following anti-CTLA4 and anti-PDL1 inhibitor treatment (Fig. 1d left, online supplementary figure 1**c**).

Synthetic long peptides (SLPs) of 21 to 27 amino acids in length have been used successfully as neoantigen vaccine approaches^20,33^. We generated SLP vaccines against 3 neoantigens (Adpgk, Reps1, Dpagt1) shown to have therapeutic efficacy in the MC38 tumor model by Yadav et al^5^. In contrast to the reported therapeutic efficacy of the SLP vaccines, the neoantigen SLP vaccines did not control tumor growth nor extended survival in MC38 tumor bearing mice compared to vehicle or CpG plus anti-CD40 adjuvant treatment alone (Fig. 1e). We then evaluated the identical long neoantigen sequences using a *Listeria monocytogenes* vector, which has been previously shown to be efficacious for tumor control when targeting long peptide sequences^29^. *Listeria monocytogenes* has been used as a cancer vaccine vector to target minimal epitopes and full-length peptides and is currently being evaluated in multiple clinical trials^34^. We generated the *Lmdda*-based Lm3 construct expressing the identical neoantigens in the aforementioned neoantigen SLP vaccine. We observed a significant reduction in tumor growth in MC38 tumor bearing mice treated with Lm3 as well as a significant increase in overall survival compared to vehicle or the ADXS-tLLO empty vector controls (Fig. 1f and 1i). To better characterize the CD8^+^ T cell response, we vaccinated mice with Adpgk SLP 27mer or minimal epitope 9mer MT peptides. The SLP vaccine against Adpgk did not increase the neoantigen-specific T cell response in the tumor compared to vehicle alone (Fig. 1g; online supplementary figure 2**a** *lower panel*). Interestingly, Lm3 was able to generate a significant intratumoral CD8^+^ T cell response to Adpgk relative to controls, which was similar to the minimal 9mer vaccine, even though Lm3 encodes for the long peptide (Fig. 1h). Collectively, these data suggest that the vaccine platform used to deliver antigen plays a crucial role in generating intratumoral immune responses and, ultimately, inhibiting tumor growth.

### Therapeutic efficacy of Listeria vectors targeting multiple neoantigens

To evaluate the potential for *Lm* immunotherapy to generate neoantigen specific immune responses against multiple neoantigens and whether these responses lead to therapeutic efficacy, we constructed the Lm20 vector targeting 20 validated neoantigens (online supplementary table 1) identified in the MC38 tumor model. The 20 neoantigens were validated using the minimal peptide screening and checkpoint inhibitor approach described in Fig. 1, with 11 neoantigens identified from 9mer peptide immunization and 9 from checkpoint inhibitor treatment (Fig. 1a-d; Fig. 2a). In addition, we generated Lm19, targeting 18 neoantigens that failed to generate a positive IFNy response by *in vivo* peptide immunization and 1 neoantigen that generated a minimal partial response following CPI treatment (Fig.1c, Fig. 1d right, and online supplementary table 1b, far right). The Lm constructs were designed as a beads-on-a-string fusion protein with adjacent neoantigen target sequences joined by linker amino acid sequences and fused to the non-hemolytic tLLO sequence required for secretion of the fusion protein into the host cell cytosol (Fig. 2b). The Lm constructs are designed to incorporate missense mutations as 21mers to allow for processing of MHC class I and class II neoantigens (Fig. 2b). We evaluated the immunogenicity and therapeutic efficacy of Lm19 and Lm20 in MC38 tumor bearing mice. Robust responses against Adpgk and Copg1, and measurable responses against >70% of targeted neoantigens were detected in the tumors following two doses of Lm20 (Fig. 2c). To our surprise, we were able to detect significant neoantigen-specific immune responses against >50% of targeted neoantigens from the tumors of mice treated with Lm19 compared to control, even though the same neoantigens failed to generate detectable T cell responses using peptide vaccination (Fig. 2d). Furthermore, both Lm19 and Lm20 treatments slowed tumor growth, and Lm20 significantly prolonged survival, with approximately 30% of Lm20 treated animals achieving a complete response (Fig. 2e). Additionally, we evaluated the capacity of ADXS-NEO to express multiple neoantigen sequences by generating a single construct targeting all 39 neoantigens included in Lm19 and Lm20. We vaccinated mice using the Lm19_20 single construct or a combination of Lm19 + Lm20. Both the Lm19_20 single construct and Lm19+Lm20 mixed vaccine significantly slowed tumor growth (online supplementary figure 3).

### Lm drives antigen-specific responses towards secondary neoantigens

To monitor the immunogenicity of individual constructs during development of the bacterial vectors for these studies, we included SIINFEKL, a validated peptide sequence known to elicit robust CD8^+^ T cell responses in C57BL/6 mice^35^. Inclusion of this sequence had the added benefit of addressing the question of whether the presence of a strong antigenic peptide sequence would result in the attenuation of T cell responses to other peptide antigens in the vaccine vector and/or a reduction in the ability to control tumors in the MC38 tumor model. To determine the effect of an immunodominant epitope, SIINFEKL, on vaccine efficacy, we immunized MC38 tumor bearing mice with *Lm* constructs expressing either tLLO alone with no target antigen (ADXS-tLLO), a immunogenic SIINFEKL model antigen not expressed in the MC38 tumor line (ADXS-tLLO-SIINFEKL), or the validated neoantigens in Lm20 without SIINFEKL (Lm20*(without SIINFEKL)) or with SIINFEKL (Lm20) included in the construct. The inclusion of the SIINFEKL sequence in Lm20 did not diminish T cell responses to other antigenic peptide sequences in the vector. Both Lm20 and Lm20* (without SIINFEKL) showed the strongest levels of tumor control (Fig. 3a, online supplementary figure 4a-b), suggesting the inclusion of SIINFEKL in the vector did not affect its capacity to control tumor growth. Strong CD8^+^ T cell responses towards multiple neoantigens were produced from the tumors and spleens of Lm20 and Lm20* (without SIINFEKL)-vaccinated mice (Fig. 3b and online supplementary figure 4c-d). Moreover, we found that ADXS-tLLO-SIINFEKL vaccinated mice generated CD8^+^ T cell responses against Adpgk, Pfkl, Chac1 and Med12 even though these antigens were not included in the vector (Fig. 3b). We also observed modest levels of tumor growth inhibition in ADXS-tLLO-SIINFEKL vaccinated mice (Fig. 3c).

**Figure 3.**
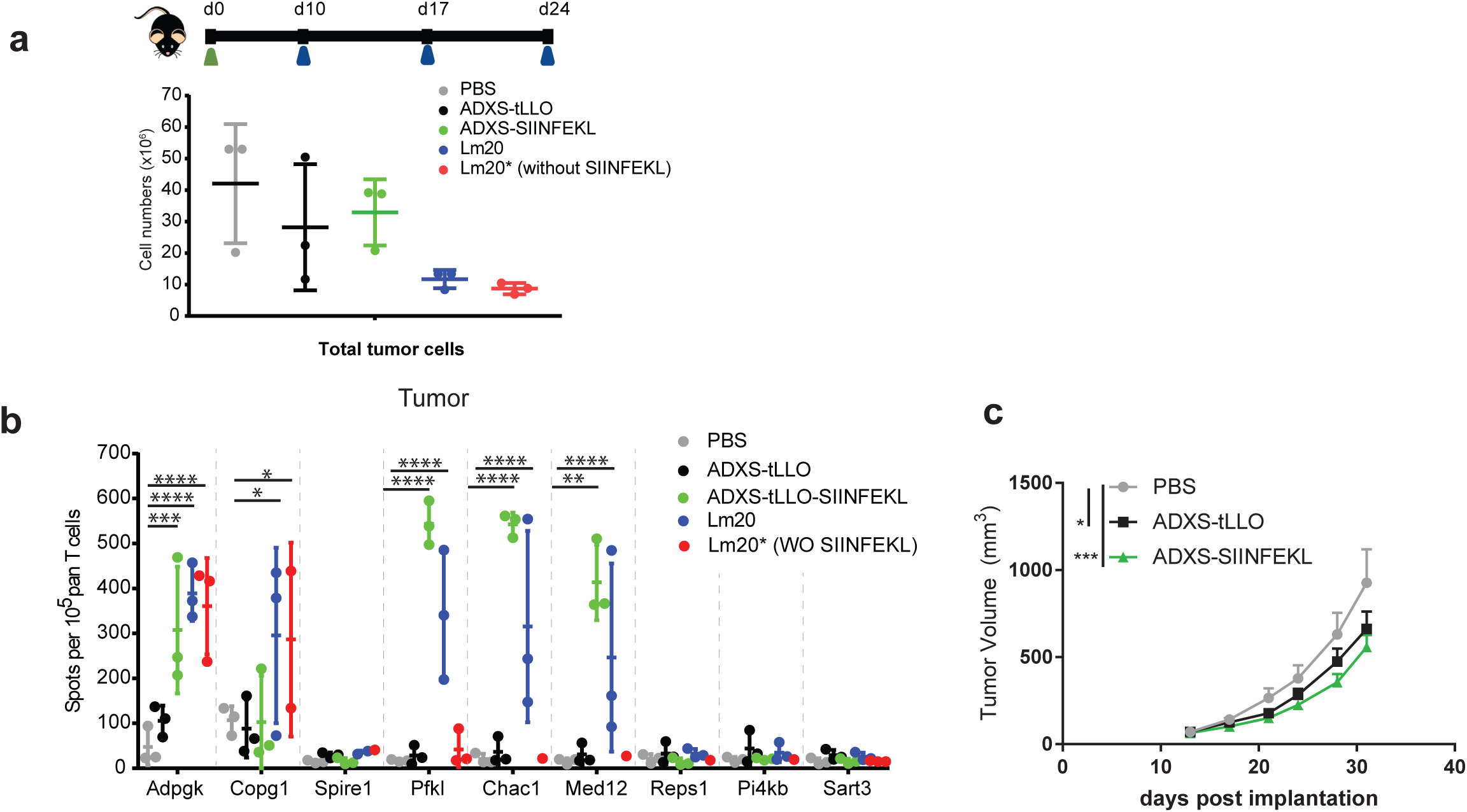
Antigen spreading by ADXS-NEO. **a**, Top, Schema of the study; bottom, total cell numbers isolated from the tumor. **b**, Representative ELISpot assay depicting the total number of IFNγ^+^ spots using ex-vivo pan T cells isolated from day 24 MC38 tumors in mice treated with either PBS or ADXS-tLLO, ADXS-tLLO-SIINFEKL, Lm20 and Lm20* (Lm20 without SIINFEKL). **c**, MC38 tumor volume kinetics in mice treated with either PBS, ADXS-tLLO (10^8^ CFU/mouse), or ADXS-SIINFEKL (10^8^ CFU/mouse). *Statistics*, ***b***, Two way ANOVA multiple comparison, *, 0.01<Adjusted P <0.05; **, 0.001< Adjusted P < 0.01; ***, 0.0001< Adjusted P <0.001; ****, Adjusted P < 0.0001. ***c***, Two way ANOVA and Tukey multiple comparison*, 0.01< P <0.05; **, 0.001< P < 0.01; ***, 0.0001< P <0.001.

Both the ADXS-tLLO-SIINFEKL and Lm20 constructs induced CD8^+^ T cell responses against the Adpgk, Pfkl, Chac1 and Med12 neoantigens in the tumor (Fig. 3b). However, T cell responses against Pfkl, Chac1, and Med12 neoantigens were abrogated in vaccinated mice with Lm20*(without SIINFEKL) vaccine when SIINFEKL was removed from the Lm20. These data suggest that the response to Pfkl, Chac1, and Med12 neoantigens in ADXS-tLLO-SIINFEKL and Lm20 vaccinated mice is dependent on the generation of a SIINFEKL response. Furthermore, CD8^+^ T cell responses against Adpgk and Copg1 from the tumor and spleen (Fig.3b and online supplementary figure 4c), and Dab2ip and Cpne1 from the spleen were maintained in Lm20*(without SIINFEKL), indicating that these responses are independent of generating a SIINFEKL response and specific to neoantigen-delivery.

Vaccination with ADXS-tLLO did not generate a significant response to neoantigens included in Lm20, although tumor growth was modestly inhibited when ADXS-tLLO was given at a higher dose (1e8 CFU/mouse), consistent with previous findings^36^ (Fig.3 b-c). However, that does not preclude the potential for antigen-specific responses against other non-targeted MC38 neoantigens. We found that mice vaccinated with ADXS-tLLO were able to generate CD8^+^ T cell responses towards neoantigens included in Lm19 (Extended Fig. 5a). The response varies between individual mice, thus we stratified ADXS-tLLO treated animals by tumor growth inhibition compared to PBS (responders vs non-responders). The ADXS-tLLO responders have a significantly higher proportion of neoantigen reactive TILs (Extended Fig. 5b). These data demonstrate the induction of resopnses against non-targeted neoantigens is an intrinsic property of *Listeria* vaccination and may be further enhanced with the inclusion of immunodominant epitopes.

### ADXS-NEO drives a pro-inflammatory tumor microenvironment

We next assessed the generation of effector CD8^+^ T cells in the TME of MC38 tumors following vaccination with ADXS-NEO (Fig. 4a and Online supplementary figure 6a). By Day 22, Lm19 and Lm20 vaccinated mice displayed early signs of tumor regression (online supplementary figure 6a). *Lm* immunization significantly increased the proportion of total TILs, CD8^+^ TILs, cytotoxic Foxp3^-^CD4^+^ TILs, γδ T cells and terminally differentiated KLRG1^lo^ CD8^+^ T cells in the tumor (Fig 4a; online supplementary figure 6b-d). The observed TME changes were not the result of antigen specific responses but, rather, were associated with immunization with the tLLO expressing Lm vector. Interestingly, ∼40% of effector TILs in untreated MC38 tumor bearing mice are of the PD-1^int/hi^ LAG-3^+^ exhausted phenotype concomitant with a reduction in the expression of the cytotoxic granule associated serine protease granzyme A (GzmA) (Fig. 4b-d). Treatment with *Lm* resulted in the emergence of a CD8^+^PD-1^lo^ effector T cell population with cytotoxic potential characterized by increased GzmA expression (Fig.4e-f). To determine if PD-1 expression is reduced in neoantigen specific T cells, we evaluated PD-1 expression on Adpgk-specific T cells. Although Adpgk-specific T cells emerge in non-vaccinated and empty vector vaccinated tumor-bearing mice, their PD-1 expression is significantly higher than mice vaccinated with constructs targeting the Adpgk neoantigen (Fig. 4g-h).

**Figure 4.**
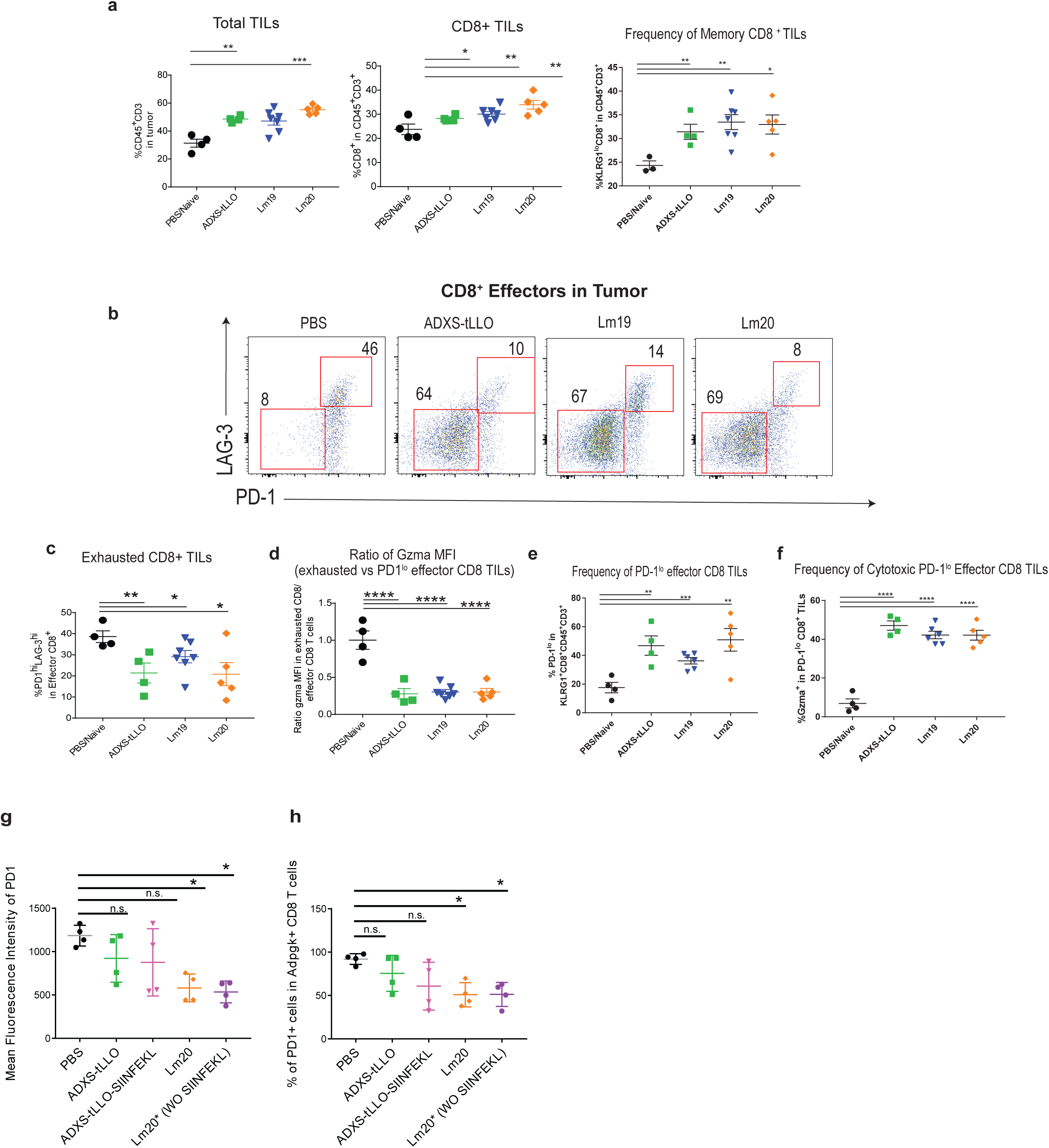
ADXS-NEO generates a robust neoantigen-specific CTL response. **a-h**, Day 22 harvested MC38 tumors, (n=4-7) in PBS, ADXS-tLLO, Lm19, or Lm20 treated mice. **a**, Left to right: Immunophenotyping of total TILs, depicting an increase in total TILs, CD8^+^ TILs, memory CD8+ TILs, **b**, Representative flow cytometric immunophenotyping CD8^+^ effectors depicting a decrease in LAG-3 and PD-1 surface expression. **c**, Frequency of exhausted (PD-1^int/hi^LAG-3^+^) CD8 TILs, **d**, Loss of Gzma expression represented by the ratio of granzyme A mean fluorescence intensity in exhausted (PD-1^int/hi^LAG-3^+^) vs effector (PD1^lo^) CD8 TILs, **e**, Frequency of effector (PD-1^lo^) CD8 TILs. **f**, Frequency of cytotoxic gzma+ effector CD8 TILs. **g**, Mean fluorescence intensity of PD-1 on Adpgk dextramer positive cells from CD8 TILs. **h**, Percentages of PD1 positive cells within Adpgk dextramer positive CD8 TILs. *Statistics*; **a-f**, Unpaired t-test, *, 0.01< P <0.05; **, 0.001< P < 0.01; ***, 0.0001< P <0.001; ****, P < 0.0001. **g**-**h**, One-way ANOVA for multiple comparisons. *, 0.01< Adjusted P <0.05; n.s., not significant.

Immune evasion in cancer is facilitated by the recruitment of regulatory T cells (Tregs), myeloid derived suppressor cells (MDSCs) and tissue associated macrophages (TAMs) into the tumor, which correlates with disease progression and decreased overall survival^37,38^. We evaluated the suppressive TME landscape in ADXS-tLLO, Lm19 and Lm20 vaccinated MC38 tumor bearing mice and found that the frequency and total number of CD4^+^CD25^hi^Foxp3^+^ intratumoral Tregs was significantly reduced (Fig. 5a-c), resulting in elevated effector CD8:Treg and effector CD4:Treg ratios (Ext. Fig. 6e-f). Likewise, the mean fluorescence intensity of Foxp3 was markedly decreased in the intratumoral Tregs compared to Lm19-or Lm20-treated mice (Fig. 5d). Foxp3 expression on a per cell basis correlates with the suppressive capacity of Tregs, suggesting that the suppressive function of Tregs in *Lm* vaccinated mice may be attenuated. The disparity of Foxp3 expression and effector:Treg ratio between ADXS-tLLO and Lm19 or Lm20 suggests an antigen-specific modulation of intratumoral Tregs. The attenuation of Tregs following *Lm* treatment is confined to the TME and was not observed in the spleen or draining lymph node (online supplementary figure 7c-d). Additionally, the TAM compartment was substantially diminished following Lm20 treatment. We detected a decrease in the quantity and proportion of TAMs and MDSCs (online supplementary figure 8a-e). Furthermore, conversion of macrophages from the pro-tumoral M2 (Arginase-I) towards the anti-tumoral M1 (iNOS) subset, characterized by both elevated iNOS production and reduced Arginase-I producing TAMs, was detected in all *Lm-*treated mice (Fig. 5e-f).

**Figure 5.**
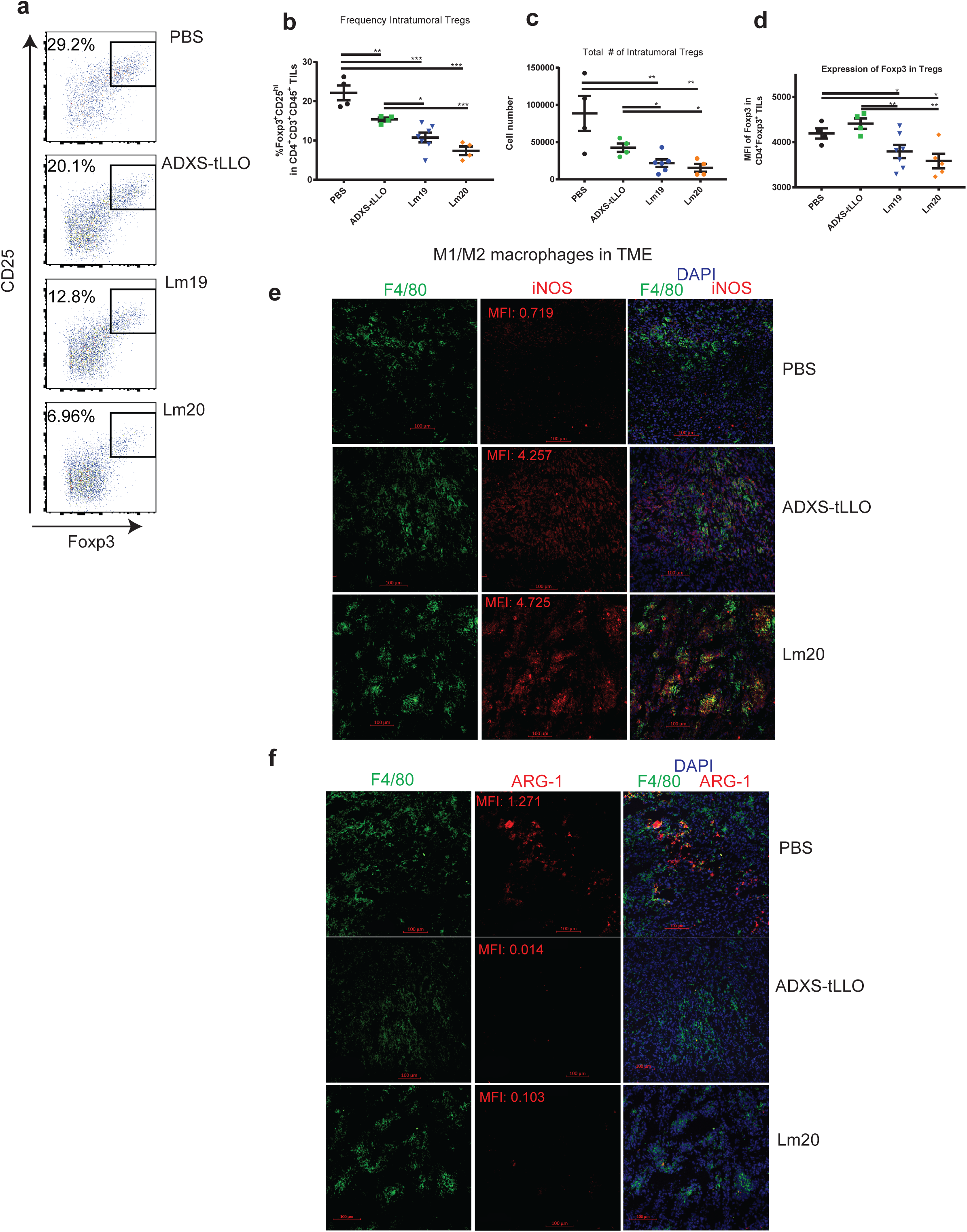
ADXS-NEO attenuates the suppressive tumor microenvironment. **a-d**, Flow cytometric phenotyping of the intratumor regulatory T cell compartment in day 22 harvested MC38 tumors in mice treated with ADXS-NEO, (n=4-7). **e-f**, Representative immunocytochemistry of day 22 harvested MC38 tumors. **a**, Gain of M1 phenotype (iNOS^+^, red) in TAMs (F4/80, green). **b**, Loss of M2 macrophages (Arg-1^+^, red; F4/80 green) following treatment with *Lm. Statistics;* Unpaired t-test, *, 0.01< P <0.05; **, 0.001< P < 0.01; ***, 0.0001< P <0.001; ****, P < 0.0001.

### ADXS-NEO therapy confers long-lasting and neoantigen-specific protective immunity

The stochastic nature of thymocyte development generates TCR repertoire diversity that is not fully heritable between individual mice and, as a result, not all mice respond equally to T cell targeted therapies^39^. We stratified responders and non-responders based on their respective tumor volume and growth curves (Fig. 6a) and evaluated T cells from the tumor and spleen. All responder mice had a higher proportion of CD45^+^ cells within the tumor, and harbored Adpgk-specific CD8^+^ T cells in the blood, spleen, and tumor, whereas non-responder mice showed Adpgk-specific CD8^+^ T cells only in the tumor (Fig. 6a,c). Responder mice generated a robust IFNγ responses towards the Adpgk and Copg1 neoantigens, as well as MC38 tumor itself, while non-responder mice did not (Fig. 6b). Additionally, responder mice had lower PD-1 expression on Adpgk-specific TILs compared to non-responders (Fig. 6d-e). Furthermore, we observed complete tumor clearance in 20-30% of vaccinated mice (Fig. 7a). Splenic T cells from tumor cleared (CR) mice contained high percentages of Adpgk-specific T cells (Fig. 7b) and generated robust CD8^+^ IFNy^+^ responses when stimulated with Adpgk or Copg1 mutated peptides or total dissociated MC38 tumor (Fig. 7c). Additionally, CR mice have a higher proportion of CD8^+^CD107a^+^ T cells, characteristic of a CTL phenotype associated with degranulation and tumoricidal capacity (Fig. 7d).

**Figure 6.**
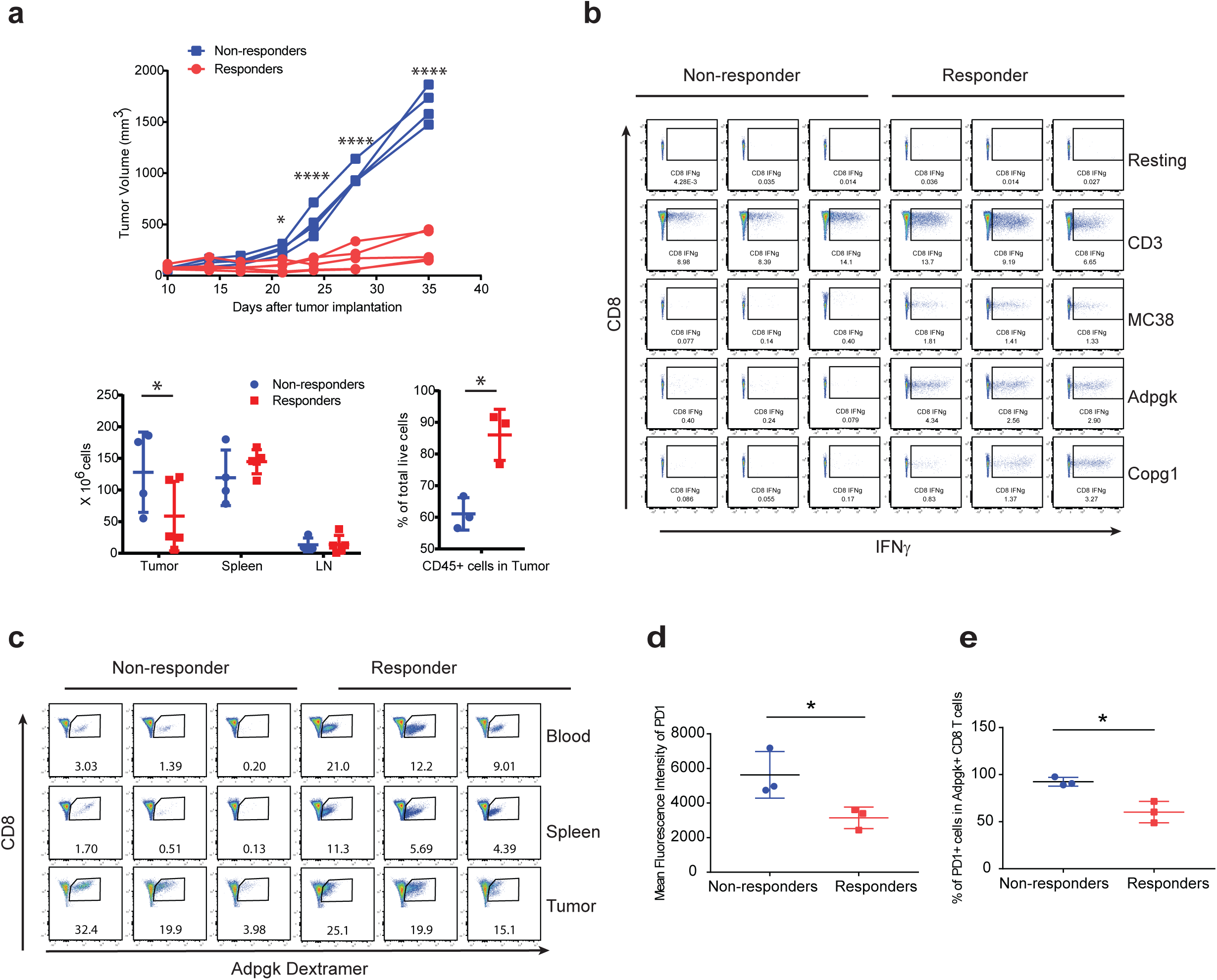
Heterogeneous immune responses observed in ADXS-NEO treated mice. **a**, Tumor growth of responder and non-responder mice immunized with Lm20*(*Top*). On day 36, mice with tumor size greater than 1200 mm^3^ were selected as “Non-responders” and mice with tumor size smaller than 500 mm^3^ were selected as “Responders.” *Bottom left*, Total cell numbers from tumor, spleen and lymph nodes from Responder and Non-responder. *Bottom right*, Percentages of CD45+ cells from tumor of responder or non-responder mice. **b**, Generation of tumor-specific IFNγ from splenic CD8 T cells from responder or non-responder mice. **c**, Neoantigen-specific CD8 T cells in the blood and spleen of responder and non-responder mice. **d**, Mean fluorescence intensity of PD1 on Adpgk dextramer positive cells from CD8 TILs of responders and non-responders. **e**, Percentages of PD1 positive cells within Adpgk dextramer positive CD8 TILs of responders and non-responders. Statistics, **a**, top and bottom left, Two-way ANOVA for multiple comparisons. *, 0.01< P <0.05; ****, P < 0.0001; bottom right, Unpaired t-test, *, P < 0.05. **d**-**e**, Unpaired t-test. *0.01< P <0.05.

**Figure 7.**
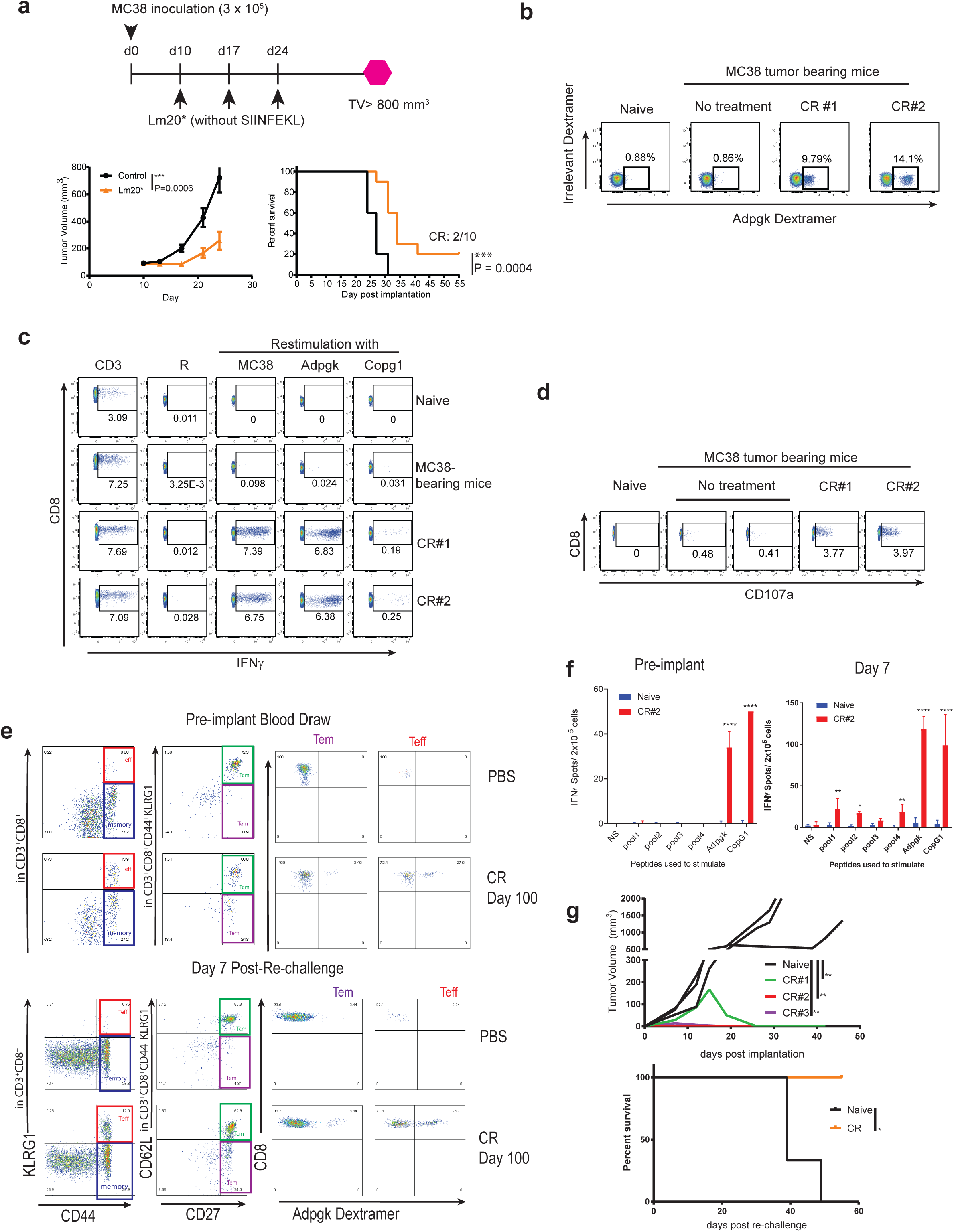
ADXS-NEO therapy confers long-lasting memory and neoantigen-specific protective immunity. **a**, Schema of the study (top). Bottom: Tumor growth(left) and survival (right) of the mice vaccinated with PBS (n=10) or Lm20*(Lm20 without SIINFEKL, n=10). **b**, Percentages of CD8 T cells recognizing mutated Adpgk assessed by Adpgk dextramer in the spleen of tumor-cleared mice 56 days after tumor implantation. CR, complete responder. **c**, Generation of IFNγ from CD8 T cells from tumor-cleared mice re-challenged with irradiated MC38 or MT peptides of Adpgk or Copg1. Spleens from tumor-cleared mice were re-challenged with minimal peptides from MT Adpgk or Copg1 or irradiated MC38. **d**, CD107a upregulation following MC38 re-challenge. CD8 T cells from tumor-cleared mice induced CD107a when re-challenged with irradiated MC38, while CD8 T cells from tumor-bearing mice failed to induce CD107a. **e-g**, Lm20 treated mice that completely cleared MC38 tumor or age-matched naïve controls were re-challenged with MC38 after being tumor free for: CR#1:198 days, CR#2: 100 days, CR#3: 216 days. **e**, flow cytometric phenotyping of circulating T cells in peripheral blood from CR#2 before tumor implantation (top panel) and 7 days post re-challenge (bottom panel). **f**, *Ex vivo* IFNγ^+^ spots circulating in the peripheral blood from CR#2 before and after MC38 re-challenge, re-stimulated with minimal peptides from Lm20 validated neoantigens in pools of 4, or 9mer of MT Adpgk and Copg1 alone. Pooling detailed in online supplementary table 2. **g**, Tumor volume kinetics and survival in MC38 re-challenged mice. *Statistics*; **a**, Tumor volume, measurement of Area Under Curve, One way ANOVA, ***, P=0.0006; Survival curve, Log-rank (Mantel-Cox) test, ***, P = 0.0004. **f**, Unpaired t-test, *, 0.01< P <0.05; **, 0.001< P < 0.01; ***, 0.0001< P <0.001; ****, P < 0.0001. **g**, Two-way ANOVA and Tukey multiple comparison*, 0.01< P <0.05; **, 0.001< P < 0.01.; Survival analysis Log-rank test *, 0.01< P <0.05.

To determine if *Lm*-generated neoantigen-specific T cell responses persist, we analyzed circulating CD8^+^ T cells in the blood of CR mice 100 and 200 days following tumor clearance. We observed a significantly higher proportion of circulating effector CD8^+^ T cells (Teff) (CD44^+^KLRg1^+^) and effector memory CD8^+^ T cells (Tem) (CD44^+^KLRG1^-^CD62L^lo^CD27^+^) in the CR mice compared to age matched naïve mice (Fig. 7e, *top*). We were also able to detect circulating Adpgk and Copg1 neoantigen-specific T cells in Lm20 treated CR mice 100 days after tumor clearance (Fig. 8e-f). Interestingly, >90% of circulating Adpgk-specific T cells from the CR mice express phenotypic markers associated with effector T cells (CD44^hi^KLRG^hi^) (Fig. 7e). We next evaluated whether long-lived memory responses were protective by re-challenging CR mice with MC38 tumor cells. One mouse completely rejected re-challenge, and the other two mice cleared the tumor by day 21. (Fig. 7g). Concurrent with protective memory, tumor re-challenge lead to the recall and expansion of circulating neoantigen-specific T cells (Fig. 7f). Although Adpgk and Copg1 -specific CD8^+^ T cells were maintained at a very low level prior to re-challenge, their responses expanded three-fold 7 days after tumor implantation (Fig. 7f).

## Discussion

Neoantigen vaccines have the potential to lead the next wave of cancer immunotherapy, as both a monotherapy and alongside checkpoint inhibitors. The success of any neoantigen cancer vaccine will be predicated on its potential to overcome two major challenges: first to generate robust neoantigen-specific effector T cell responses with tumoricidal potential and second to convert a “cold” tumor microenvironment (TME) to “hot” by engaging both the innate and adaptive arms of the immune system. Here we establish, using a pre-clinical murine tumor model, that ADXS-NEO, a genetically engineered *Listeria monocytogenes* cancer vaccine, successfully overcomes these challenges. With the capacity to target upwards of 40 potential neoantigens per construct, we demonstrate that ADXS-NEO elicits robust T cell responses towards neoantigens in the MC38 tumor model. Moreover, because ADXS-NEO is a live attenuated bacterial vector, systemic administration of the *Lm* vaccine drives pro-inflammatory innate and adaptive immunity and converts the “cold” TME to “hot,” by attenuating intratumoral Tregs, MDSCs, and M2 TAMs and generating effector αβ/γδ tumor infiltrating T cells and M1 macrophages. Altogether, these data identify ADXS-NEO as a promising personalized immunotherapy platform with a high capacity to drive effector T cell responses against multiple neoantigens while simultaneously engaging innate immune cells, ultimately producing robust and long-lived immunity towards tumor derived neoantigens.

Recent advances in whole genome sequencing have enabled the rapid identification of non-synonymous mutations from a relatively small tumor biopsy, making a truly personalized cancer vaccine possible^21^. The quantity of non-synonymous mutations (NSMs) are highly varied between tumor types, from potentially thousands in melanoma, lung and bladder cancers to single digits in prostate, glioblastoma, and various leukemias ^21^. While a greater number of NSMs correlates with more neoantigen targets and an increased potential for anti-tumor immunity ^40^, large numbers of NSMs also pose technical obstacles for the design and manufacture of cancer vaccines. The choice of which antigenic targets to include in a cancer vaccine is critical due to inherent limitations in vector capacity. This makes antigenic prediction both a necessary step, as well as a potential limitation in the production of personalized cancer vaccines. Various algorithms have emerged with the goal of predicting the binding affinity of a potential neoantigen for a given MHC molecule. Even with improvements in our ability to predict peptide-MHC binding affinity, accurate neoantigen selection is further limited by our ability to predict TCR binding to the pMHC complex, and ultimately the capacity to generate a tumoricidal effector T cell response. Therefore, it is critical that any system used to deliver tumor-specific neoantigens for the purpose of eliciting anti-tumor T cell immunity be as efficient as possible with respect to the number of neoantigens successfully targeted.

For example, as shown in Figure 1, we observed that only 9 of 37 predicted tested neoantigens generated significant immune responses in all mice immunized using a minimal peptide + adjuvant approach. Furthermore, we showed that validated neoantigens were unable to control tumor growth when administered as a synthetic long peptide vaccine. Yet when these same validated neoantigens were delivered using the ADXS-NEO vector they were able to generate T cell responses that slowed MC38 tumor growth. Moreover, the ADXS-NEO Lm19 construct, targeting non-validated neoantigens that failed to elicit a strong IFNγ response via peptide vaccination, not only generated neoantigen specific responses, but controlled tumor growth and extended overall survival. These findings highlight the disparity between the immunogenic potential of different vaccine delivery systems^41^, a potential that is further complicated by the route of immunization^42^.

Considering the limited effectiveness of the currently available predictive neoantigen algorithms, strategies for maximizing the effectiveness of personalized vaccines are paramount. One such strategy is to empirically validate potential neoantigens by screening patient PBMCs for neoantigen reactivity. However, this approach will miss neoantigens that fail to prime responses in the patient, either due to sub-optimal conditions present in “cold” TME’s or the lack of sufficient antigenic protein for cross-presentation and priming by dendritic cells. Furthermore, empirical validation of individual neoantigens adds time to an already time constrained manufacturing process, making it of limited utility in treating patients in more advanced stages of disease. In the studies presented here, we demonstrate that neoantigen validation by minimal peptide immunization or by checkpoint inhibitor blockade, was not indicative of neoantigen immunogenicity when delivered by an intracellular *Lm*-based vaccine, as Lm19 treated mice were able to generate T cells expressing IFNγ in response to non-validated neoantigens.

Another strategy for overcoming the limitations of predictive neoantigen selection is to simultaneously target all or the majority of NSMs, thereby de-emphasizing the importance of accurate neoantigen prediction by increasing the quantity of potential neoantigens delivered by the vaccine (increasing the “shots on goal”). While certain high mutational load tumor types have hundreds of potentially targetable NSMs, most tumor types have fewer than 10 somatic mutations per Mb, which corresponds to ∼150 NSMs, and many have fewer than 1 somatic mutation per Mb^3^. Building on the “shots on goal” strategy for neoantigen vaccine design, we demonstrate that ADXS-NEO has the capacity to deliver upwards of 40 neoantigens per individual construct, with a potential to simultaneously immunize with multiple constructs while maintaining tumor control efficacy in the MC38 model (online supplementary figure 3). As NGS technology improves and multi-biopsy approaches emerge that may increase the quantity of variant calls^43^, cancer vaccines, such as ADXS-NEO, with the capacity to deliver many neoantigens simultaneously provides a platform that can address the unpredictability of neoantigen identification.

A pre-clinical study using an mRNA-based neoantigen vaccine reported that CD4^+^ T cells are required for tumor control, while CD8^+^ T cells play a minimal role^44^. Furthermore, early clinical data from an mRNA-based personalized cancer vaccine trial in melanoma found that the majority of vaccine induced antigen-specific T cell responses were CD4^+^ T cell rather than CD8^+^ T cell mediated, despite the use of an algorithm designed to predict MHC class I binding peptides for the identification of neoepitope sequences to include in the vaccine constructs^19^. In contrast to these findings, we showed that ADXS-NEO induced a robust neoantigen-specific CD8^+^ T cell response, likely due to the inherent bias towards MHC class I antigen presentation and generation of CD8^+^ T cell responses associated with *Lm* infection ^31^. However, ADXS-NEO also generates effector CD4^+^ helper T cell responses, allowing for the establishment of long-lasting memory and protection against tumoral relapse. Interestingly, we found circulating neoantigen-specific effector and effector memory T cells in the blood of ADXS-NEO treated mice more than 100 days after tumor clearance. Importantly, ADXS-NEO treatment preferentially generates a less exhausted PD-1 low population of neoantigen-specific TILs, and these PD-1 low TILs were shown to correlate with tumor control. Although neoantigen-specific TILs were detected in non-vaccinated and empty-vector vaccinated tumor-bearing mice, these TILs were PD-1 high. It is possible that PD-1 high T cells may be pre-existing, a phenomenon that may be related to T cell activation resulting from cross-presentation in non-vaccinated mice compared to direct MHC class I presentation in Lm20 treated mice. The skewing towards PD-1 low TILs and their overall contribution to tumor control, ability to establish a long-lived memory response, and overall survival warrants further investigation.

Our findings demonstrate that an *Lm*-based personalized cancer vaccine can generate a strong effector and effector memory CD8^+^ T cell response able to control tumor growth and provide long-lasting protection in mice. With a high capacity to target many neoantigens and the ability to maintain a pro-inflammatory tumor microenvironment, ADXS-NEO has the potential to overcome many of the obstacles impeding the success of personalized cancer vaccines. However, the overall efficacy of PCVs has yet to be established in the clinic. ADXS-NEO, along with many other PCV platforms, are currently being evaluated in phase I clinical trials.

## Declarations

### Ethics approval and consent to participate

All data are generated using mice and there is no participant in this study.

### Consent for publication

All authors agree with publication. The material presented in this submission has not been previously reported and is not under consideration for publication elsewhere.

### Availability of data and material

All data are included in the manuscript and supplementary information.

### Authors’ contributions

BC conceived, designed and performed experiments, analyzed data, and wrote the manuscript; OP performed experiments and analyzed data; DP, EF, XJ, CM, KR, NT, ZC, BVL, JL, XL, JD, KC, SL, JZ, PM and DOV performed experiments and analyzed data; DB analyzed data; SMH, JAJ, and RP contributed the design of the study and analyzed the data; and HP and MFP conceived and designed the study, supervised the research, analyzed data, and wrote the manuscript.

## Acknowledgements

We thank Jing Qing and Wenjun Ouyang for reading the manuscript and providing intellectual input.

